## Supplementary figures and images for "Improving gut virome comparisons using predicted phage host information"

### Supplementary Figure 1

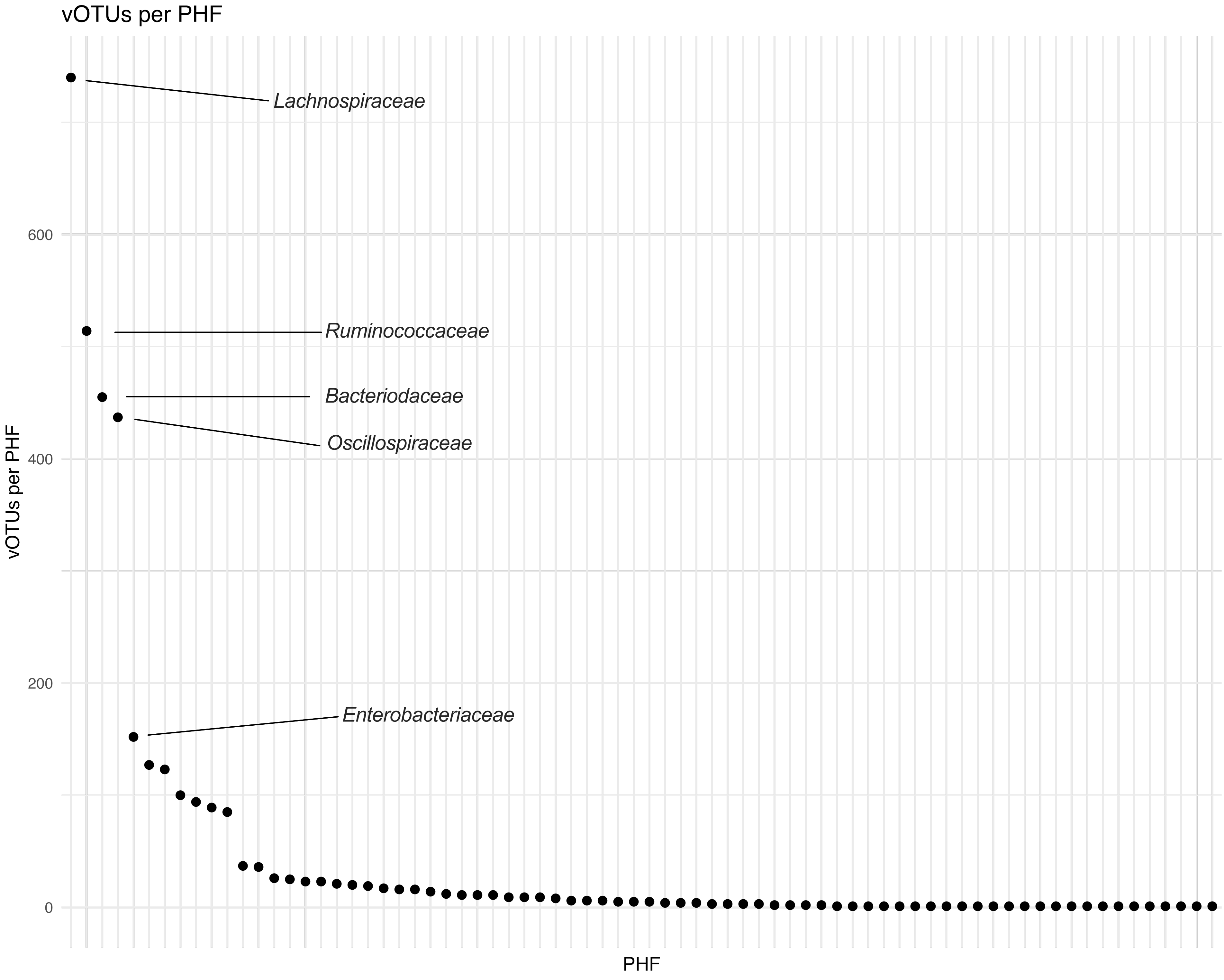

### Supplementary Figure 2

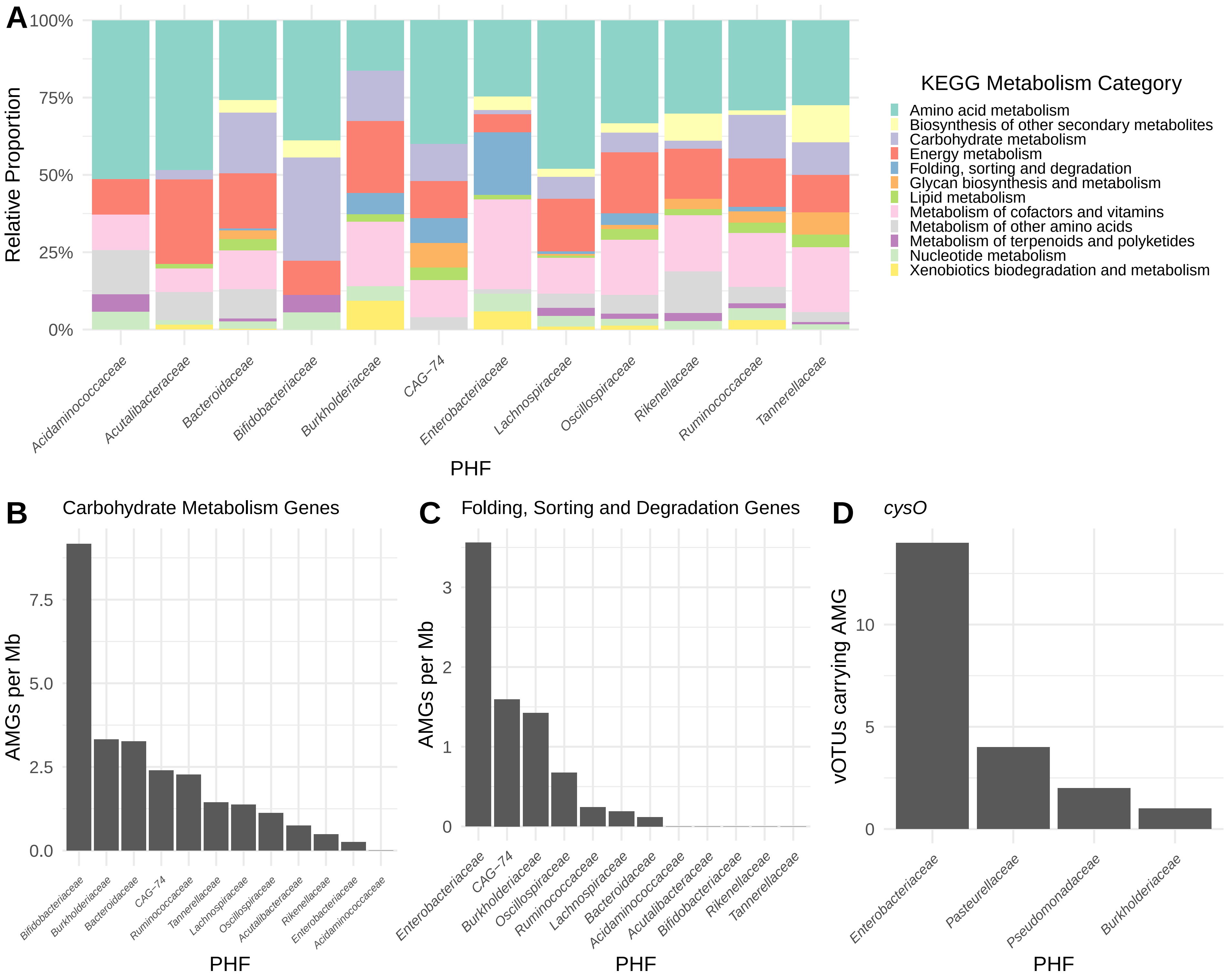

### Supplementary Figure 3

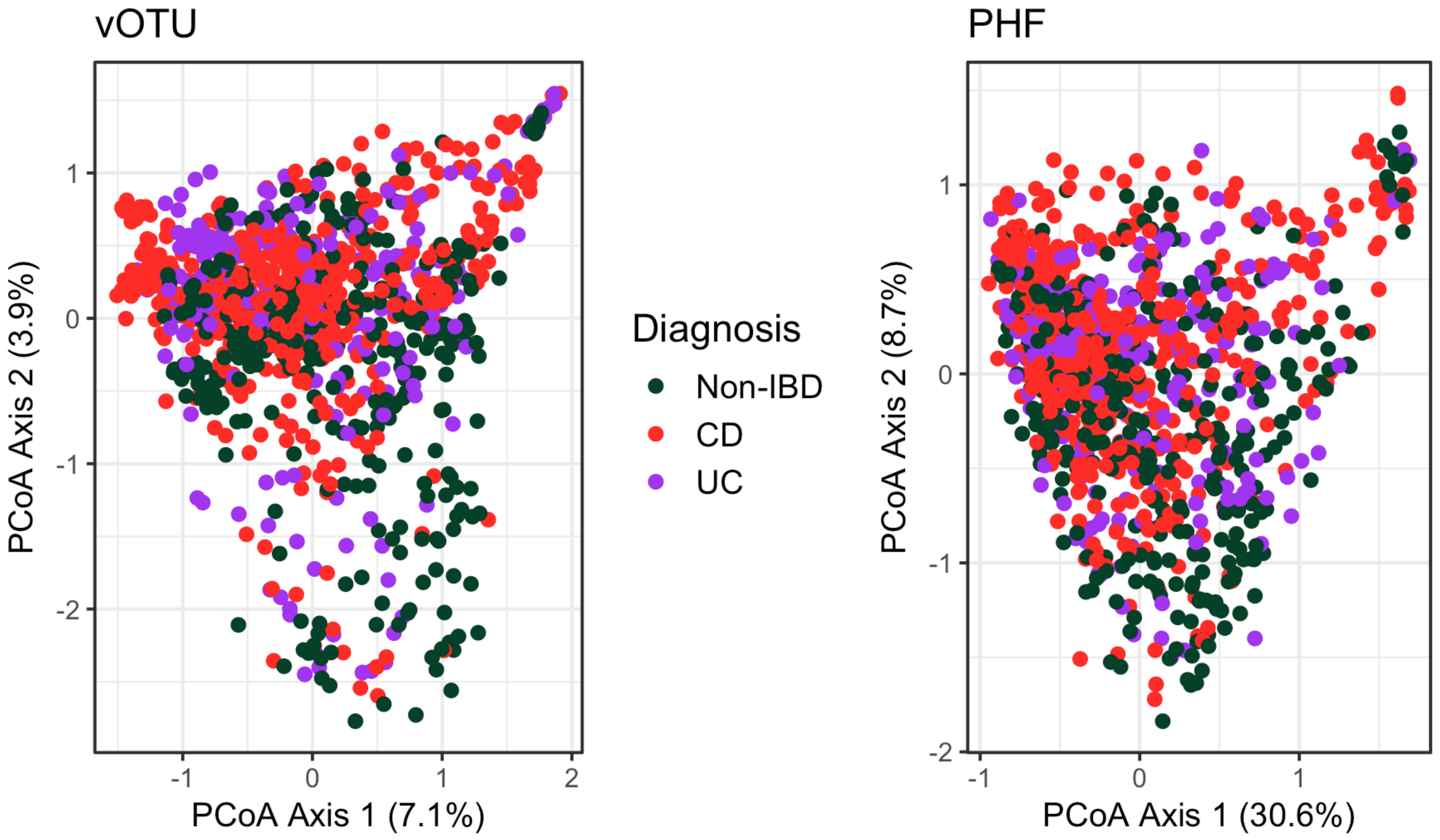
